# Deletion of can1/cat1 genes and expression of a dominant any1 mutation establish an effective canavanine selection in fission yeast

**DOI:** 10.1101/2022.03.11.484000

**Authors:** Anissia Ait Saada, Alex B. Costa, Kirill S. Lobachev

## Abstract

Positive and counter-selectable markers have been successfully integrated as a part of numerous genetic assays in many model organisms. In this study, we investigate the mechanism of resistance to arginine analog canavanine and its applicability for genetic selection in *Schizosaccharomyces pombe*. Deletion of both arginine permease genes *cat1* and *can1* provides strong drug resistance, while the single *can1* deletion does not have impact on canavanine resistance. Surprisingly, the widely used *can1-1* allele does not match to the *can1* gene but rather corresponds to the *any1-523C>T* allele. The strong canavanine-resistance conferred by this allele arises from an inability to deposit basic amino acid transporters on the cellular membrane. *any1-523C>T* leads to reduced post-translational modifications of Any1 regulated by the Tor2 kinase. We also demonstrate that *any1-523C>T* is a dominate allele. Our results uncover the mechanisms of canavanine-resistance in fission yeast and open the opportunity of using *cat1, can1* and *any1* mutant alleles in genetic assays.

## Introduction

Counter-selectable markers are widely used as genetic tools to isolate cells that experienced a mutation leading to the loss/inactivation of the marker. Counter selection in *S. pombe* relies mostly on the use 5-FOA (5-fluoroorotic acid) and, to a lesser extent, FudR (5-fluoro-2’-deoxyuridine) to select for mutations in *ura4* and *TK* (herpes virus thymidine kinase) genes, respectively. In budding yeast, counterselection on canavanine, a toxic arginine analog, has been successfully integrated as a part of numerous genetic assays (e.g. (Myung et al., 2001; Yang et al., 2008; Zhang et al., 2013). This approach takes advantage of the *CAN1* gene that encodes for an arginine permease and confers resistance to canavanine upon its loss of function. In contrast, canavanine resistance is not part of the repertoire of markers in *S. pombe*, although arginine permeases and mutants conferring canavanine resistance are known. So far, counterselection on canavanine has been limited to plasmid shuffling using the *S. cerevisiae CAN1* gene to complement the *S. pombe can1-1* mutation (Ekwall and Ruusala, 1991; Paluh and Clayton, 1996).

Canavanine cytotoxicity is due to its structural similarity with arginine. Canavanine is a substrate for tRNA arginyl synthase and is incorporated into nascent proteins instead of arginine. The resulting proteins harbor an altered structure and function, which leads to cells death. Therefore, in cells that do not discriminate between arginine and its toxic analog, canavanine resistance is strictly correlated with a defect in arginine uptake. The uptake of arginine, as well as other amino acids and nucleobases, is tightly regulated by the localization and expression of their cognate transporters. This regulation is sensitive to the nutritional environment and involves different signaling pathways like Tsc/Rheb in fission yeast and target of rapamycin (TOR) in both fission and budding yeasts (Aspuria and Tamanoi, 2008; Liu et al., 2015; Ma et al., 2013; MacGurn et al., 2011).

In *S. cerevisiae*, arginine can be transported by three amino acid permeases: Can1, Gap1, and Alp1, with Can1 being the main transporter (Zhang et al., 2016). In *S. pombe*, two arginine permeases have been identified: Can1 (which is non-orthologous to Can1 in *S. cerevisiae*) and Cat1 (ScAlp1 homolog). In contrast to budding yeast, Cat1 is the main arginine permease (Aspuria and Tamanoi, 2008). This is surprising since a notoriously used allele, *can1-1*, that confers strong resistance to canavanine was thought to correspond to a mutation in the *can1* gene (Fantes and Creanor, 1984). While little is known about Can1, the control of amino acid uptake via the Cat1 permease is well studied in fission yeast. Cellular localization of Cat1 defines the rate of arginine/canavanine uptake, and the Tsc1-Tsc2 complex is required for proper Cat1 distribution on the plasma membrane (Aspuria and Tamanoi, 2008). Accordingly, cells defective for the Tsc/Rheb pathway are canavanine resistant and show an intracellular mislocalizaton of Cat1. Delocalization of Cat1 from the plasma membrane occurs through endocytosis of the permease by Any1 (a β-arrestin-like protein) in cooperation with the E3-ubiquitin ligase Pub1 (Nakashima et al., 2014). It has been found that over-expression of Any1 confers resistance to canavanine, while the absence of Any1 or its ubiquitination by Pub1 confers sensitivity. In addition to ubiquitination, Any1 is subject to another post-translational modification (PTM), the role and the nature of which remain unknown (Nakase et al., 2013; Nakashima et al., 2014).

In an attempt to make canavanine counter-selection a reliable marker in *S. pombe*, we undertook the characterization of the *can1-1* allele. Surprisingly, we found that *can1* deletion does not confer resistance to canavanine but rather enhances resistance in the absence of Cat1. Whole-genome sequencing of the *can1-1* strain revealed a point mutation in *any1* and no mutations in *can1*. We found that this single mutation, *any1-523C>T*, is necessary and sufficient to reproduce the *can1-1* phenotype, *i.e*. strong canavanine resistance and Cat1 internalization. At the protein level, cells expressing *any1-523C>T* do not show an over-expression of Any1 but rather a strong diminution of both PTMs. These results reveal the importance of the unknown PTM since its absence is enough to compensate for the lack of Any1 ubiquitination. Last, we report that both Any1 PTMs are differentially regulated by the TORC1 pathway. By revealing the nature of the mutation in *can1-1*, our data add a new layer to our comprehension of the mechanisms of canavanine resistance and also the regulation of arginine uptake in *S. pombe*.

## Material and methods

### Yeast strains and growth media

The *S. pombe* strains used in this study are listed in Table 1. Gene tagging was performed by classical molecular genetics techniques (Moreno et al., 1991). Gene deletion was performed using the *delitto perfetto* approach, initially developed in budding yeast (Storici and Resnick, 2006). Briefly, the pKL421 plasmid was constructed carrying a CORE cassette containing the *kanMX6* and *ura4* markers. The CORE cassette was inserted at the target locus and selected for on YES media containing G418. The CORE cassette was then replaced by transformation with an oligomer containing homology to the flanking regions of the target locus. Transformants were selected on 5-FOA containing media. Uracil auxotroph and G418 sensitive clones were selected and tested for the absence of the CORE cassette and target gene by PCR. Cat1 at C terminus was tagged with yeGFP and *any1* was tagged at C-terminus with 6 copies of HA tag using pYM25 and pYM16 (both Euroscarf) as templates for PCR, correspondingly. Oligonucleotides used in *delitto perfetto* and gene tagging are available upon request. For cell sensitivity to canavanine, serially diluted cell suspensions were spotted on EMM media supplemented with adenine, leucine and uracil and containing 5 g/L of ammonium chloride (if not otherwise stated). L-Canavanine (Sigma C9758) was added directly to the media at the indicated concentrations.

**Table 1:**
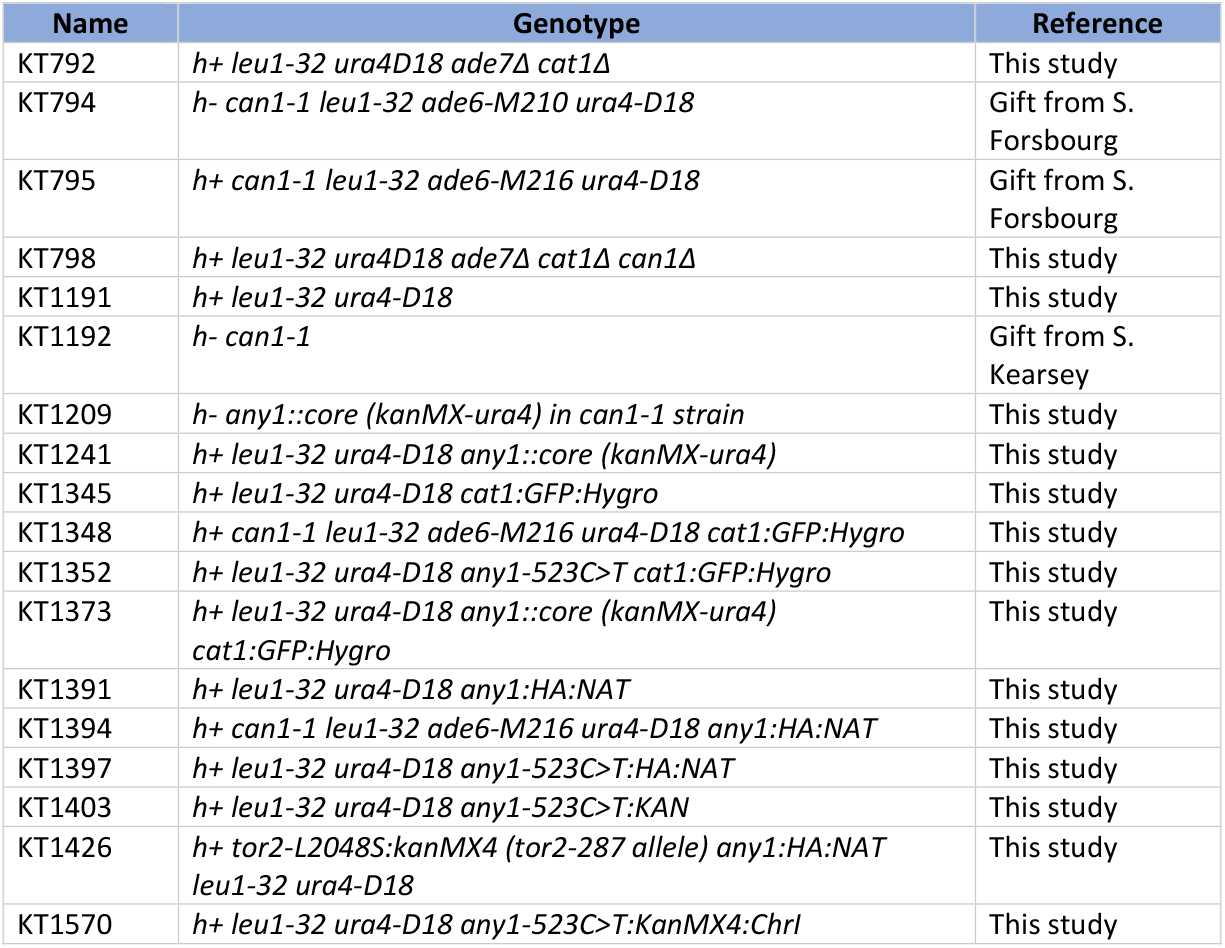
Strains used in this study.

### Genome-wide sequencing

Two *can1-1* strains were obtained from two independent labs and used for sequencing. Whole-genome sequencing was performed by Psomagen (Rockville, MD) using Illumina TruSeq DNA Nano 350 bp library prep and an Illumina NovaSeq 6000. Reads were aligned to the *S. Pombe* reference genome (Wood et al., 2002) using the Burrows-Wheeler Aligner (Li and Durbin 2009), Picard (http://broadinstitute.github.io/picard), SAMtools (Li and Durbin, 2009), and Genome Analysis Toolkit v3.8 (Auwera and O’Connor, 2020). Variants were called against the reference genome using Mutect2 (Auwera and O’Connor, 2020) and referenced against genes known to affect canavanine sensitivity (Harris et al., 2021).

### Whole protein extraction and western blot

Exponentially growing cells inoculated in YES were arrested with 0.1% sodium azide. 1×10^8^ cells were collected to perform the following protein extraction. The cell pellet was suspended and washed in 1 ml stop buffer (50 mM NaF, 10 mM NaN3 in PBS 1X) and then washed in 1 ml of 20% trichloroacetic acid (TCA). Pelleted cells were resuspended in 200 μl 20% TCA and glass beads (Sigma G8772) were added to each tube. The cell walls were mechanically broken using a FastPrep-24 bead beating homogenizer (MP-Biomedicals) with the following program: 6000 rpm, 3 rounds of 30 sec ON with a one-minute interval between each round on ice. 400 μl 5% TCA was added to the cell lysate and cell lysate was recovered without the beads in a new tube. The cell lysate was spun at 13k rpm for 5 min and the pellet was resuspended in 200 μl TCA buffer (1x SDS loading buffer, 0.2M Tris-HCl pH 8). The samples were denatured by boiling at 95°C for 5 min followed by a brief centrifugation prior to western blot analysis. Proteins were resolved by 4-20% SDS-PAGE gel and then transferred onto a PVDF membrane. The membrane was probed with anti-HA (1:2000, Sigma H6908) or anti-H3 (1:5000, Novus NB500-171) antibodies and the proteins were revealed by chemiluminescence with horseradish-peroxidase-conjugated sheep anti-rabbit IgG (cytiva).

### Live cell imaging

Cells were inoculated in filtered EMM-NH4Cl medium supplemented with adenine, leucine and uracil. Cells were prepared for microscopy as described in (Pietrobon et al., 2014). Exponentially growing cells were washed in fresh supplemented EMM media and a 2 μl drop was deposited on the well of a microscope slide (Thermo Scientific, ER-201B-CE24) covered with a layer of 1.4% agarose in filtered EMM. Images were acquired with the Zeiss LSM 700A confocal microscope.

## Results

### The Can1 permease plays a minor role in arginine/canavanine uptake

In fission yeast, expression of the *can1-1* allele and deletion of *cat1* are known to confer resistance to canavanine. The nature of the *can1-1* allele remained unknown since its discovery in 1984 but, as its name indicates, *can1-1* was thought to correspond to a mutation in the *can1* gene. The allele is, indeed, referred to as defective *can1* gene in literature. However, the *can1* gene is not used as a counter-selectable marker in *S. pombe*. We reasoned that if *can1-1* encodes for a defective Can1 arginine permease, the deletion of *can1* should confer resistance to canavanine to the same extent as *can1-1*. We found that deletion of *can1* does not provide substantial resistance to canavanine compared to *cat1* deletion or *can1-1* strains (Figure 1). Although deletion of *can1* slightly enhances canavanine resistance of the *cat1* mutant, the fact remains that the *can1-1* strain displays the strongest canavanine resistance. These results suggest that Can1 plays a minor role in arginine and canavanine uptake compared to Cat1. Importantly, these results indicate that the *can1-1* phenotype does not flow from a mutation in the *can1* gene.

**Figure 1:**
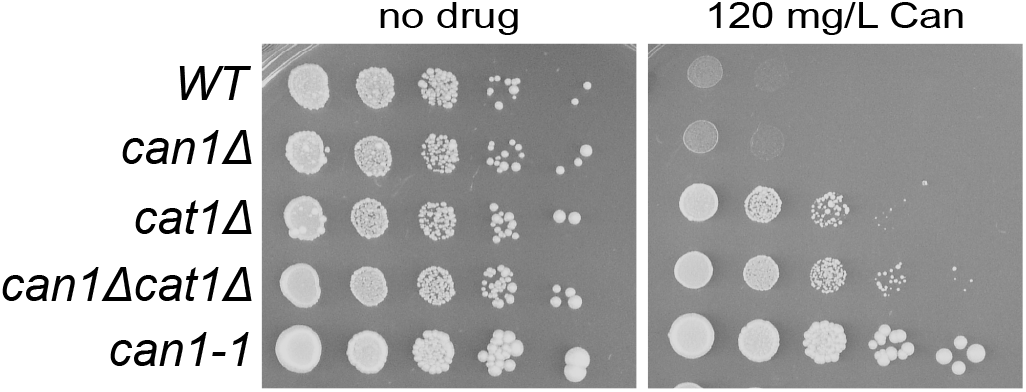
Canavanine resistance in arginine permease mutants. Serial 10-fold dilutions of indicated strains spotted on synthetic media (EMM) lacking arginine and containing 5 g/L of ammonium chloride in the absence or presence of canavanine (Can). The medium was supplemented with adenine, leucine and uracil.

### The *can1-1* allele corresponds to a mutation in the *any1* gene

Chromosome mapping of *can1-1* showed that the corresponding gene is located on chromosome II and is linked to *tps13* (a centromere-linked marker), which is 35 map units distant (Fantes and Creanor, 1984). Confusingly, the *can1* gene resides around this location but is not involved in canavanine resistance (Figure 1). A simple but pertinent hypothesis explaining the *can1-1* phenotype is the involvement of another gene located in close proximity to *can1*. In order to determine the genetic basis of canavanine resistance attributed to the *can1-1* allele, we performed whole-genome of two strains from Dr. S. Forsburg laboratory FY254 (h-*can1-1 leu1-32 ade6-M210 ura4D18*) and FY261 (h+ *can1-1 leu1-32 ade6-M216 ura4D18*). During the analysis of sequencing data, we focused our attention on mutations (i) located around the deduced *can1-1* location and/or (ii) present in genes involved in canavanine resistance. One particular mutation stood out because it presented both criteria. Indeed, the *can1-1* strain showed a point mutation (523C>T) in *any1* gene which (i) is located only 8.4 kb away from *can1* and (ii) encodes for an arrestin-related endocytic adaptor involved in the regulation of amino acid transporters like Cat1 and Aat1. Knock out of *any1* confers an extreme sensitivity to canavanine, whereas its overexpression confers resistance to canavanine (Nakase et al., 2013; Nakashima et al., 2014).

To verify that this mutation is sufficient to recapitulate the canavanine resistance observed in the *can1-1* strain, we recreated the *any1-523C>T* in our strain background and deleted *any1* in the *can1-1* strain. We observed that cells expressing *any1-523C>T* show the same resistance to canavanine as the original *can1-1* strain. In addition, the deletion of *any1* in *can1-1* not only suppresses canavanine resistance but also exacerbates sensitivity to the drug like observed in the *WT*strain lacking *any1*. Last, the replacement of *any1* gene in the *can1-1* strain with the *WT any1* sequence suppresses canavanine resistance in this strain and shows the same sensitivity as the *WT* strain. These results show that canavanine resistance in the *can1-1* strain is not due to a defective arginine permease, like so far assumed, but rather due to a gain of function mutation in the regulator of arginine transporters, Any1. Although we found that the mutation responsible for canavanine resistance in the strain isolated by Fantes and Creanor resides in *any1* gene, we will keep referring to the strain as *can1-1* to distinguish it from the strain harboring *any1-523C>T* that we recreated in our strain background.

Fantes and Creanor noticed that nitrogen source impacts growth of *can1-1* in the presence of canavanine. The *can1-1* strain presented a better resistance to the drug when cells were grown on ammonium versus glutamate (two nitrogen sources used in *S. pombe* media). It is also known that nitrogen starvation leads to an abundance of amino acid transporters at the plasma membrane, which ensures a greater nutrient uptake (MacGurn et al., 2011; Nakashima et al., 2014). Therefore, canavanine sensitivity increases upon nitrogen stress because of a greater uptake of the drug. We tested if the canavanine resistance of the *can1-1* strain is impacted by the amount of available nitrogen in the media. We used 0.5, 2 and 5 g/L of ammonium chloride, the highest concentration being the standard concentration used in *S. pombe* media. Expectedly, we observed that the extent of canavanine sensitivity in the *WT* strain is greater in the presence of limiting amounts of ammonium (Figure 2B). Also, the canavanine resistance of cells deleted for both of the arginine permeases, Can1 and Cat1, is lessened by limiting nitrogen availability. Interestingly, decreasing the amount of nitrogen did not have a drastic impact on canavanine resistance in the *can1-1* strain as well as *any1-523C>T* (Figure 2B). This suggests that nitrogen starvation does not lead to an accumulation of amino acid transporters at the cell surface in cells expressing *any1-523C>T*. We noticed though that *any1-523C>T* presents a slightly slower growth compared to the *can1-1* strain. This might be explained by the differences in the genetic background of the two strains.

**Figure 2:**
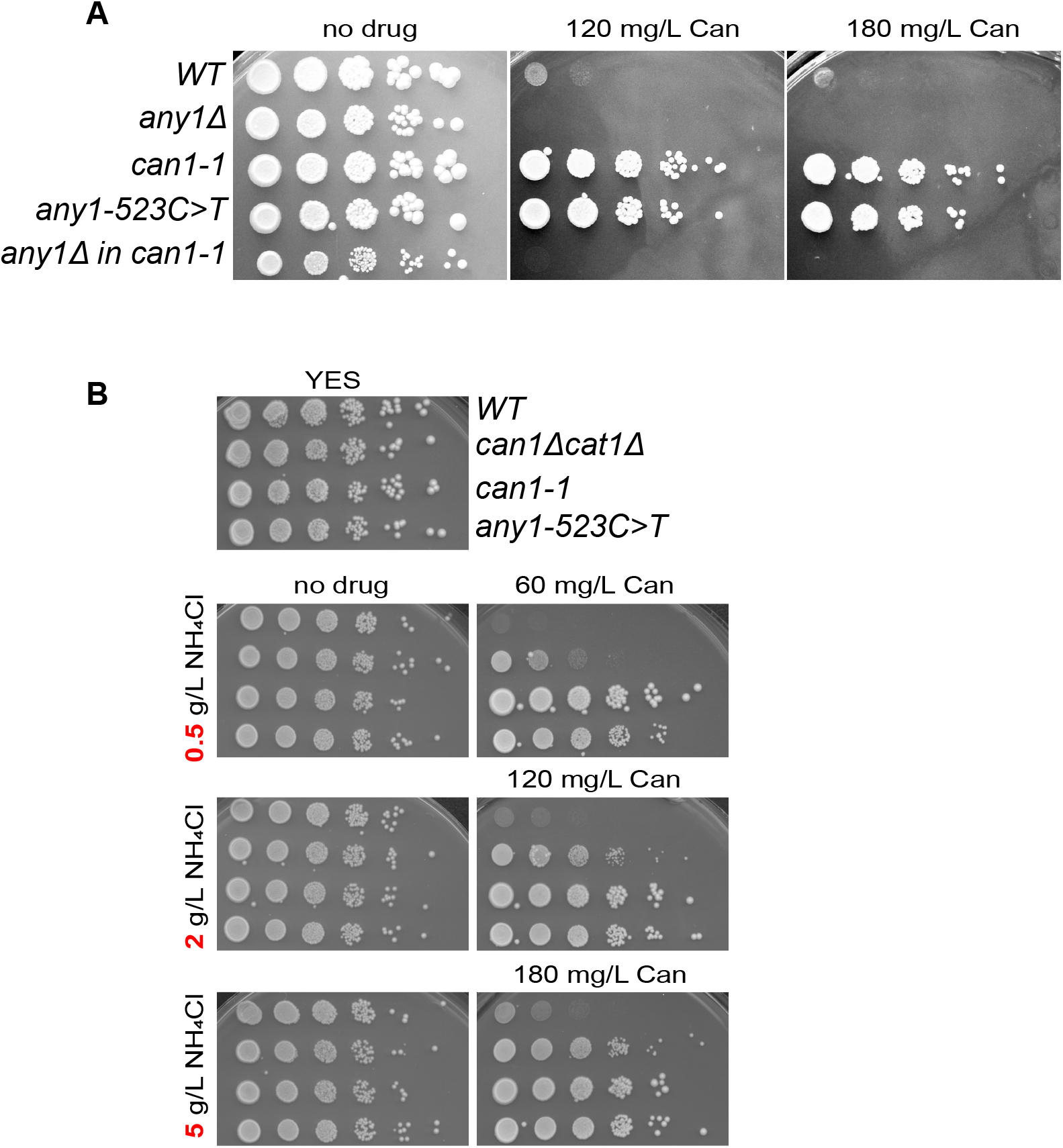
A point mutation in *any1* is responsible for canavanine resistance in the *can1-1* strain. **A.** Serial 10-fold dilutions of indicated strains spotted on synthetic media lacking arginine and containing 5 g/L of ammonium chloride in the absence or presence of canavanine. **B.** Serial 10-fold dilutions of indicated strains spotted on YES or synthetic medium lacking arginine and containing the indicated amount of ammonium chloride and canavanine (Can).

### *any1-523C>T* is a dominant allele in haploid strains

The fact that deletion of *any1* leads to an acute canavanine sensitivity whereas *any1-523C>T* leads to a clear-cut resistance suggests that *any1-523C>T* is a gain-of-function mutation. In most cases, gain-of-function mutations are dominant or semi-dominant. To test whether *any1-523C>T* is dominant, we created a heterozygote strain where *any1^+^* is expressed from its endogenous locus and *any1-523C>T* is inserted in the left arm of chromosome I. Therefore, both a *WT* and a mutated copy of Any1 are expressed in the haploid strain. We found that the heterozygote (*any1^+^/any1-523C>T*) strain shows the same capability to grow on canavanine-containing media as *can1-1* and *any1-523C>T* even under nitrogen starvation conditions (Figure 3). This confirms that *any1-523C>T* is a dominant allele and also offers a great opportunity to use this allele as a selectable marker without the need to delete the endogenous gene.

**Figure 3:**
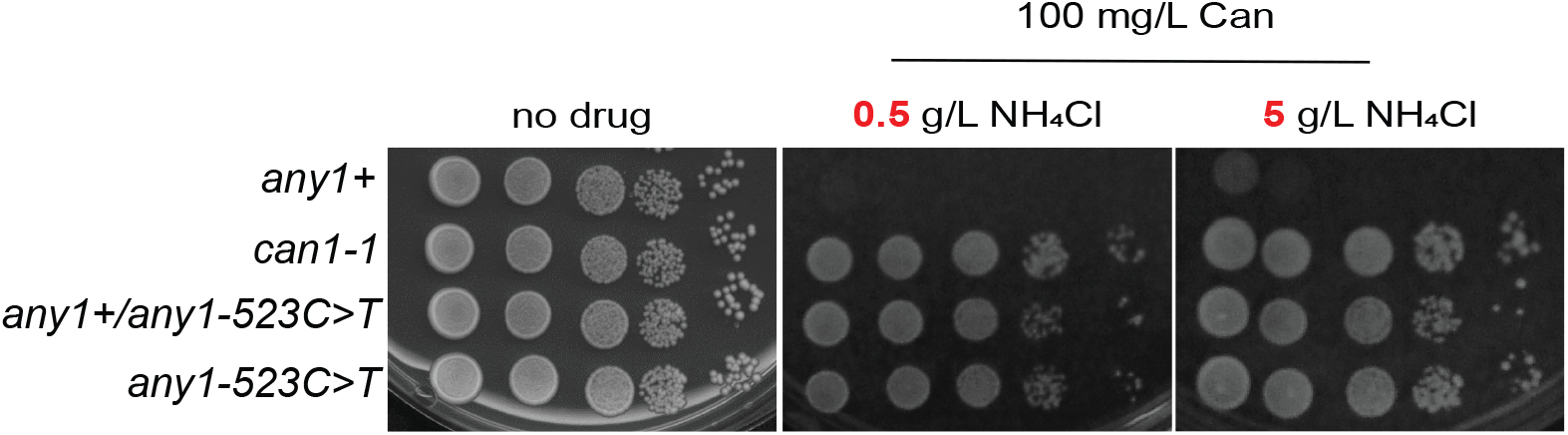
*any1-523C>T* is a dominant allele. Serial 10-fold dilutions of indicated strains spotted on synthetic media lacking arginine and containing the indicated amount of ammonium chloride and canavanine (Can). The strains are haploid and express either one copy of *any1* (*any1+, can1-1, any1-523C>T*) or two copies (*any1+/any1-523C>T*). The heterozygous strain contains *any1+* in its endogenous location and *any1-523C>T* inserted into the left arm of chromosome I.

### Expression of Any1^R175C^ changes the post-translational modification pattern

To further understand the mechanism of canavanine resistance in *can1-1*, we analyzed the effect of the 523C>T mutation, which induces a change of amino acid 175 from an arginine to a cysteine. This mutation is located in the arrestin motif of the protein (Figure 4A). Nakashima et al. (2014) showed that *any1^+^* overexpression confers canavanine resistance and that ubiquitination of Any1 at the K263 residue by Pub1 is required for Cat1 endocytosis. Indeed, deletion of *pub1* and expression of Any1^K263R^ confer sensitivity to canavanine to the same extent as *any1* deletion.

**Figure 4:**
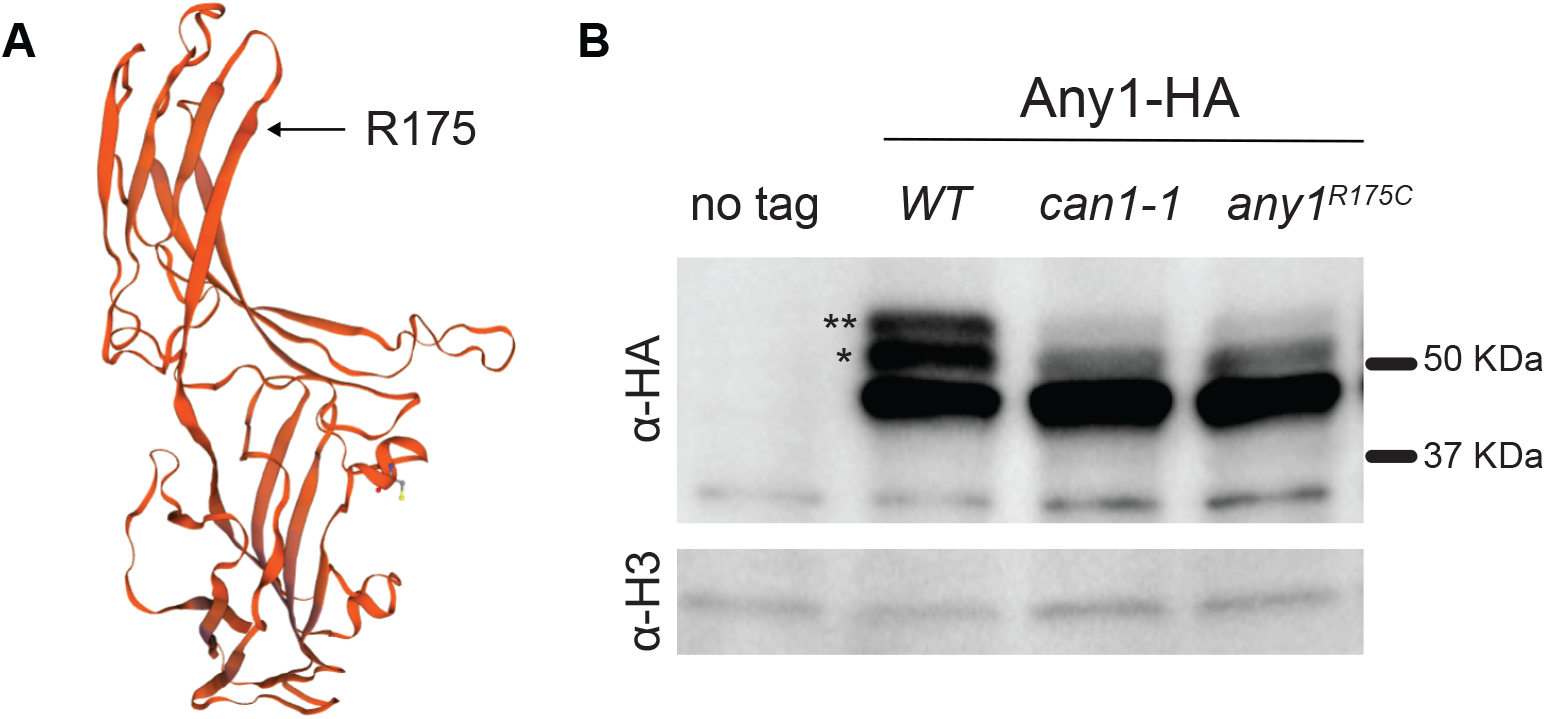
Effect of the *any1-523C>T* mutation at the protein level. **A.** Automated protein structure prediction using homology modeling (SWISSMODEL). Expression of *any1-523C>T* leads an arginine to cysteine substitution at position 175. **B.** Expression of *WT* and mutated Any1 (*can1-1* and Any1-R175C) analyzed by immunoblotting the endogenously tagged Any1 with anti-HA. Histone H3 is shown as a loading control. The asterisks correspond to the post-translationally modified forms of Any1, where * corresponds to an unknown PTM and ** corresponds to the Pub1-mediated ubiquitinated form. Both PTMs are strongly diminished in cells expressing Any1-R175C.

We tagged the endogenous *any1* gene in a *WT* strain and in cells expressing *any1-523C>T* (either in *can1-1* or in our strain background). As expected, on a western blot, we observed three distinct bands in the *WT* strain (Nakashima et al., 2014): a main band and two electrophoretic mobility shifts where the upper band (*) corresponds to the ubiquitinated Any1 and the middle band (**) corresponds to an unknown PTM (Figure 4B). The protein status in *can1-1* and *any1-523C>T* was unexpected in terms of correlation with canavanine resistance. First, expression of Any1^R175C^ does not lead to a higher level of the protein that would mimic *any1+* overexpression. Second, Any1^R175C^ does not show an increase but rather a strong decrease in its ubiquitination level. Third, the unknown PTM, the role of which is also unknown, is diminished in Any1^R175C^. This result shows that Pub1-mediated ubiquitination of Any1 is not required for canavanine resistance in cells expressing Any1^R175C^. It also suggests a potential role for the unknown PTM, loss of which may bypass the need for Any1 ubiquitination.

### Expression of Any1^R175C^ leads to a massive Cat1 endocytosis

Any1 is involved in the regulation of amino acid transporters (Nakase et al., 2013). Under nutrient-rich conditions, it is notably responsible for storage of plasma membrane transporters, like Cat1 and Aat1, in the Golgi apparatus (Nakase et al., 2013; Nakashima et al., 2014). The fact that canavanine resistance comes from the inability of the cells to import the toxic drug implies that the gain of function mutation *any1-523C>T* leads to a massive internalization of amino-acid transporters. To test this hypothesis, we tagged the native arginine permease Cat1 with GFP in order to track its cellular localization by fluorescent microscopy (Figure 5). Analysis of GFP signal across the cell (white bar on the left panel) showed two major peaks in *WT* and *any1Δ* at the cellular periphery. This means that in these strains, Cat1 is mainly localized in the cellular membrane, thus explaining canavanine sensitivity. However, we found that in *any1-523C>T* and the original *can1-1* strains Cat1-GFP localization is mostly intracellular. This explains why the strains are very resistant to canavanine: the transporters responsible for canavanine import are excluded from the plasma membrane.

**Figure 5:**
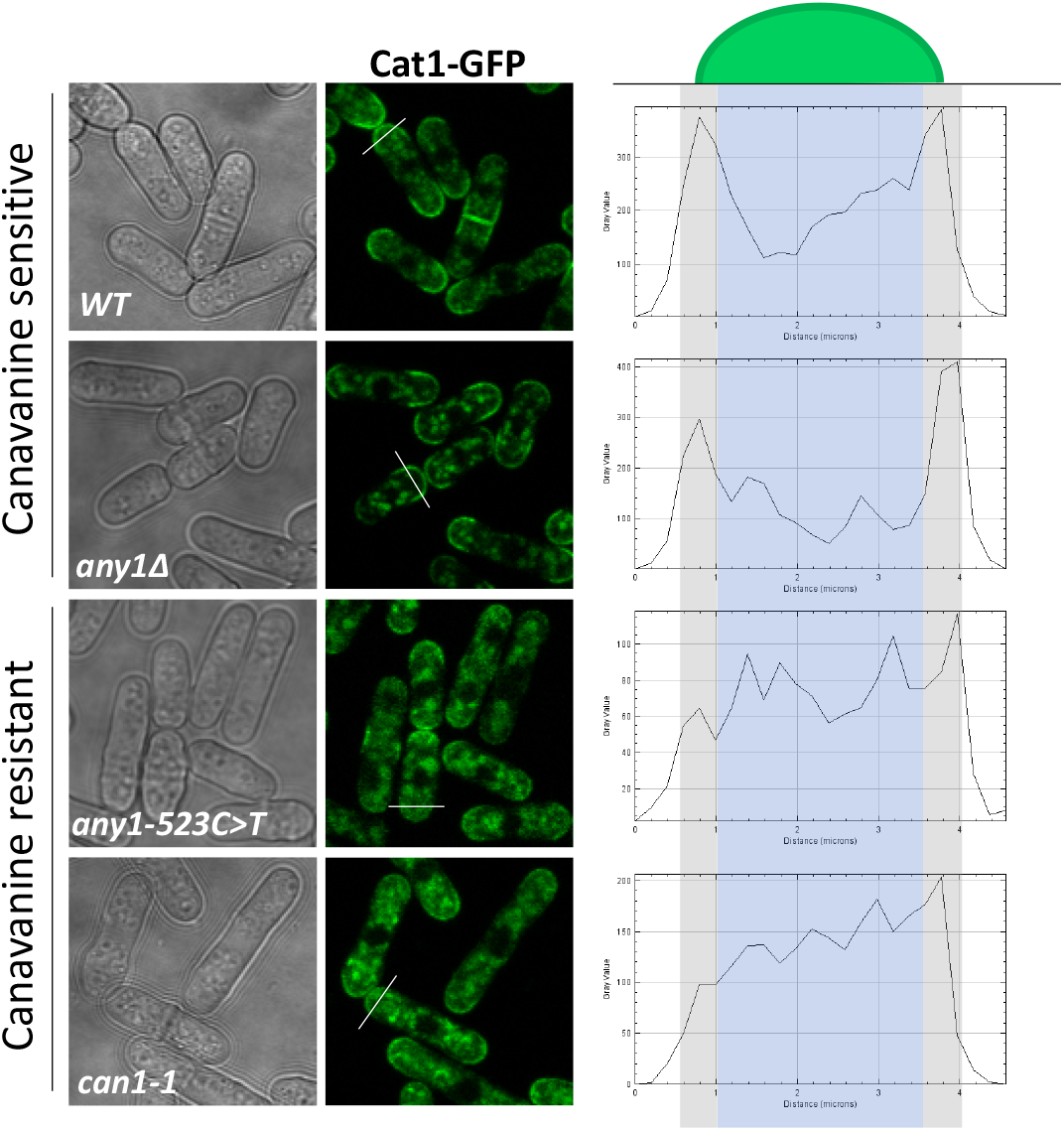
Expression of Any1-R175C leads to Cat1 internalization. Cellular localization of Cat1-GFP in indicated strains. Left panel: cells expressing Cat1-GFP observed under a fluorescent microscope. Right panel: intensity of GFP signal across the cell (white bar on the left panel).

### Any1 PTMs are differentially regulated by the TORC1 pathway

TORC1 and TORC2 are known to regulate arginine and leucine uptake. Tor2 is a protein kinase that is part of the TORC1 complex. It has been shown that activating mutations in Tor2 render cells more sensitive to canavanine whereas inactivating mutations confer canavanine resistance (Ma et al., 2013; Murai et al., 2009). The kinase defective and thermosensitive *tor2* allele (*tor2-287*, (Hayashi et al., 2007)) and, paradoxically, *tor2^+^* overexpression confer canavanine resistance and a defect in leucine uptake (Ma et al., 2013; Weisman et al., 2007). In *S. cerevisiae*, the TORC1 complex stimulates Can1-mediated arginine uptake via Npr1-mediated phosphorylation of Art1 (Any1 ortholog) (MacGurn et al., 2011). The fact that *tor2-287* confers canavanine resistance and phosphorylation of the Any1 homolog in budding yeast negatively regulates its function prompted us to verify the PTM status of Any1 in cells expressing *tor2-287*. The results show that Any1 exhibits the same PTM pattern in both *WT* and *tor2-287* at the permissive temperature (26°C) but not at the restrictive temperature (32°C) (Figure 6). Indeed, both Any1 PTMs were differentially affected in the *tor2-287* strain. The ubiquitinated form of Any1 dramatically increased whereas the unknown PTM underwent a drastic decrease. This shows the unknown Any1 PTM is positively regulated by Tor2 and suggests that its loss is linked to canavanine resistance in *any1-523C>T/can1-1*.

**Figure 6:**
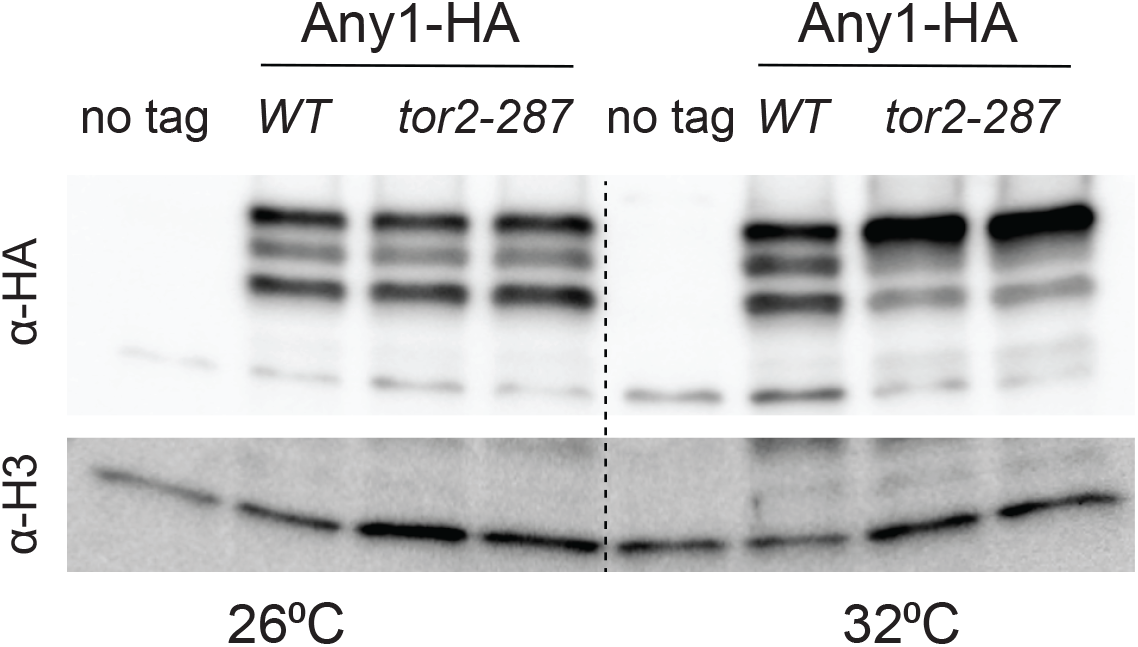
Any1 PTMs are regulated by the TORC1 pathway. Expression of Any1 in cells expressing *WT tor2* or the thermosensitive allele *tor2-287* (Tor2-L2048S). The cells were grown to exponential phase and then incubated at permissive (26°C) or restrictive (32°C) temperatures. Any1 was probed by immunoblotting with anti-HA. Histone H3 is shown as a loading control.

## Discussion

Canavanine is a valuable drug in both fundamental and clinical research. First, counterselection using canavanine has been instrumental in developing genetic assays in budding yeast (but not in fission yeast). Second, canavanine offers a great therapeutic potential in the treatment of several cancers, including breast cancer (Bence and Crooks, 2003; Karatsai et al., 2020; Nurcahyanti and Wink, 2017). Therefore, understanding the mechanism of canavanine resistance in eukaryotes is important for developing new tools in model organisms and predicting success of drug treatment in medicine.

Canavanine resistance is observed in cells lacking either an arginine permease or factors involved in the positive regulation of amino acid/arginine uptake, like the Tsc1-Tsc2 complex. *can1-1* was isolated more than three decades ago and it had been assumed that the mutation resided in the *can1* gene. In this paper we not only show that *can1-1* corresponds to a mutation in *any1*, but also deletion of *can1* does not confer canavanine resistance like expected when an arginine permease is deleted. However, deletion of *can1* and *cat1* confers substantial canavanine resistance, even upon nitrogen starvation (Figure 1 and 2B). Yet, canavanine resistance of *can1-1/any1-523C>T* is more profound than the deletion of both arginine permeases, suggesting that one or several other amino acid transporters regulated by Any1 are involved in arginine uptake. Accordingly, the mechanism by which canavanine resistance occurs in *can1-1/any1-523C>T* is via and excessive internalization of amino acid transporters, including Cat1 (Figure 5). It has been shown that *CAN1* from budding yeast functionally complements *can1-1* in fission yeast (Ekwall and Ruusala, 1991). Since *can1-1* corresponds to *any1-523C>T*, it is likely that Any1 does not control the cellular localization of the budding yeast Can1, thus explaining the complementation. It is interesting to note that *CAN1* orthologs listed in the *S. pombe* database (https://www.pombase.org) correspond to several genes encoding amino acid transporters like *cat1, aat1*, and *isp5* but not *can1*. Indeed, the *can1* ortholog corresponds to *VHC1* in budding yeast, a vacuolar membrane cation-chloride cotransporter. Although deletion of *can1* slightly enhances canavanine resistance in the absence of *cat1*, we suspect that Can1 in fission yeast is not a true arginine permease.

Besides arginine and lysine (basic amino acids), Any1 is involved in leucine uptake via the regulation of Aat1 (Cherkasova et al., 2021). Usually, mutants like *tsc2* that show canavanine resistance show also a defect in leucine uptake, which prevents the mutant cells from growing on synthetic media supplemented with leucine (Ma et al., 2013; Matsumoto et al., 2002). The strains used in this study are auxotrophic for leucine because they carry the *leu1-32* mutation. It was, therefore, expected that the canavanine resistance of *can1-1*/*any1-523C>T* would be accompanied by a growth defect on EMM media supplemented with leucine (no drug condition on figures). However, this was not the case (Figure 1 and 2). This suggests that the mutation *any1-523C>T* specifically affects the basic amino acid transporters involved in arginine trafficking and does not limit Aat1 leucine uptake. Sychrova et al. isolated a UV-induced mutant resistant to thialysine (a toxic analog of lysine) named *tlh1*. This mutant was genetically mapped at the same locus as *can1-1* and showed resistance to canavanine without affecting leucine or glutamate uptake (Sychrová et al., 1992). We predict that a follow-up analysis of the *tlh1* mutant described by Sychrova et al., will show that the allele corresponds to a mutation in *any1*.

The fact that *any1-523C>T* is a dominant mutation that elicits canavanine resistance without affecting leucine uptake, makes this allele a newly defined selectable maker that can be used in fission yeast genetics regardless of leucine auxotrophy. It should be kept in mind that cells prototrophic for basic amino-acids (arginine, lysine and histidine) should be used. In addition of positive selection using any1-523C>T, we propose the use of *can1* and *cat1* for negative selection on canavanine. This will offer the opportunity to develop canavanine-based genetic assays in fission yeast.

It is known that Tor2 regulates amino-acids transporters both positively and negatively via a complex signaling pathway. The *ts* allele *tor2-287* that confers canavanine resistance shows an increase in *cat1* expression but no apparent massive endocytosis of the permease (Ma et al., 2013). Nakase et al. found that Tor2 activity is not required for the formation of the Any1-Pub1 complex. Here, we show that Tor2 regulates Any1 at the posttranslational level. The increase in Any1 ubiquitination in *tor2-287* suggests that Tor2 modulates the interaction between Any1 and Pub1. This constitutes the first evidence that TORC1 regulates the function of the Any1-Pub1 complex. One common feature between *tor2-287* and *can1-1/any1-523C>T*, besides canavanine resistance, is a decrease in the level of the unknown PTM of Any1. It is possible that the unknown PTM corresponds to a Tor2-mediated Any1 phosphorylation. However, it has been shown recently that Any1 is subjected to an increase in phosphorylation at the T12 residue upon inhibition of the TOR pathway (Mak et al., 2021). The fact that both *tor2-287* and *can1-1/any1-523C>T* show canavanine resistance but the Any1 PTM pattern is different shows that the mechanism of canavanine resistance differs but, in both cases, involves Any1.

In humans, the Tuberous Sclerosis Complex is a pathology caused by mutations in the TSC1-TSC2 complex. Mutations in one of the tumor suppressor genes, *TSC1* or *TSC2*, lead to an aberrant hyperactivation of the mammalian TOR pathway. Yet, treatments of TSC with mTOR inhibitors are not always efficient (Luo et al., 2022). In the light of our results, it is conceivable that the defects observed in *TSC1* or *TSC2* deficient cells may involve an arrestin-mediated mechanism.

## Acknowledgements

We are very thankful to Dr. S.A.E. Lambert for insightful discussion of the results and providing several strains and oligos. We also thank Dr. S. Forsburg and Dr. S. Kearsey for providing strains containing the *can1-1* allele. Funding of this work was provided by NIH grant R01GM129119 and a Mayent Rothschild fellowship from Curie Institute (to K.S.L.).

## Author contributions

Conceptualization: A.A.S. and K.S.L.; Methodology: A.A.S., A.B.C. and K.S.L.; Investigation: A.A.S., A.B.C., and K.S.L; Writing – Original Draft: A.A.S. and K.S.L.; Writing – Review & Editing: A.A.S., A.B.C. and K.S.L.; Funding Acquisition: K.S.L.; Resources: K.S.L.; Supervision: K.S.L.; Project Administration: K.S.L.

## References

Aspuria, P.J., and Tamanoi, F. (2008). The Tsc/Rheb signaling pathway controls basic amino acid uptake via the Cat1 permease in fission yeast. Mol Genet Genomics 279, 441–450.

Auwera, G.A.V.d., and O’Connor, B.D. (2020). Genomics in the Cloud: Using Docker, GATK, and WDL in Terra (1st Edition).

Bence, A.K., and Crooks, P.A. (2003). The mechanism of L-canavanine cytotoxicity: arginyl tRNA synthetase as a novel target for anticancer drug discovery. J Enzyme Inhib Med Chem 18, 383–394.

Cherkasova, V., Iben, J.R., Pridham, K.J., Kessler, A.C., and Maraia, R.J. (2021). The leucine-NH4+ uptake regulator Any1 limits growth as part of a general amino acid control response to loss of La protein by fission yeast. PLoS One 16, e0253494.

Ekwall, K., and Ruusala, T. (1991). Budding yeast CAN1 gene as a selection marker in fission yeast. Nucleic Acids Res 19, 1150.

Fantes, P.A., and Creanor, J. (1984). Canavanine resistance and the mechanism of arginine uptake in the fission yeast Schizosaccharomyces pombe. J Gen Microbiol 130, 3265–3273.

Hayashi, T., Hatanaka, M., Nagao, K., Nakaseko, Y., Kanoh, J., Kokubu, A., Ebe, M., and Yanagida, M. (2007). Rapamycin sensitivity of the Schizosaccharomyces pombe tor2 mutant and organization of two highly phosphorylated TOR complexes by specific and common subunits. Genes Cells 12, 1357–1370.

Karatsai, O., Shliaha, P., Jensen, O.N., Stasyk, O., and Rędowicz, M.J. (2020). Combinatory Treatment of Canavanine and Arginine Deprivation Efficiently Targets Human Glioblastoma Cells via Pleiotropic Mechanisms. Cells 9.

Li, H., and Durbin, R. (2009). Fast and accurate short read alignment with Burrows-Wheeler transform. Bioinformatics 25, 1754–1760.

Liu, Q., Ma, Y., Zhou, X., and Furuyashiki, T. (2015). Constitutive Tor2 Activity Promotes Retention of the Amino Acid Transporter Agp3 at Trans-Golgi/Endosomes in Fission Yeast. PLoS One 10, e0139045.

Luo, C., Ye, W.R., Shi, W., Yin, P., Chen, C., He, Y.B., Chen, M.F., Zu, X.B., and Cai, Y. (2022). Perfect match: mTOR inhibitors and tuberous sclerosis complex. Orphanet J Rare Dis 17, 106.

Ma, N., Liu, Q., Zhang, L., Henske, E.P., and Ma, Y. (2013). TORC1 signaling is governed by two negative regulators in fission yeast. Genetics 195, 457–468.

MacGurn, J.A., Hsu, P.C., Smolka, M.B., and Emr, S.D. (2011). TORC1 regulates endocytosis via Npr1-mediated phosphoinhibition of a ubiquitin ligase adaptor. Cell 147, 1104–1117.

Mak, T., Jones, A.W., and Nurse, P. (2021). The TOR-dependent phosphoproteome and regulation of cellular protein synthesis. Embo j 40, e107911.

Matsumoto, S., Bandyopadhyay, A., Kwiatkowski, D.J., Maitra, U., and Matsumoto, T. (2002). Role of the Tsc1-Tsc2 complex in signaling and transport across the cell membrane in the fission yeast Schizosaccharomyces pombe. Genetics 161, 1053–1063.

Moreno, S., Klar, A., and Nurse, P. (1991). Molecular genetic analysis of fission yeast Schizosaccharomyces pombe. Methods Enzymol 194, 795–823.

Murai, T., Nakase, Y., Fukuda, K., Chikashige, Y., Tsutsumi, C., Hiraoka, Y., and Matsumoto, T. (2009). Distinctive responses to nitrogen starvation in the dominant active mutants of the fission yeast Rheb GTPase. Genetics 183, 517–527.

Myung, K., Datta, A., and Kolodner, R.D. (2001). Suppression of spontaneous chromosomal rearrangements by S phase checkpoint functions in Saccharomyces cerevisiae. Cell 104, 397–408.

Nakase, Y., Nakase, M., Kashiwazaki, J., Murai, T., Otsubo, Y., Mabuchi, I., Yamamoto, M., Takegawa, K., and Matsumoto, T. (2013). The fission yeast β-arrestin-like protein Any1 is involved in TSC-Rheb signaling and the regulation of amino acid transporters. J Cell Sci 126, 3972–3981.

Nakashima, A., Kamada, S., Tamanoi, F., and Kikkawa, U. (2014). Fission yeast arrestin-related trafficking adaptor, Arn1/Any1, is ubiquitinated by Pub1 E3 ligase and regulates endocytosis of Cat1 amino acid transporter. Biol Open 3, 542–552.

Nurcahyanti, A.D., and Wink, M. (2017). L-Canavanine Potentiates Cytotoxicity of Chemotherapeutic Drugs in Human Breast Cancer Cells. Anticancer Agents Med Chem 17, 206–211.

Paluh, J.L., and Clayton, D.A. (1996). Mutational analysis of the gene for Schizosaccharomyces pombe RNase MRP RNA, mrp1, using plasmid shuffle by counterselection on canavanine. Yeast 12, 1393–1405.

Pietrobon, V., Fréon, K., Hardy, J., Costes, A., Iraqui, I., Ochsenbein, F., and Lambert, S.A. (2014). The chromatin assembly factor 1 promotes Rad51-dependent template switches at replication forks by counteracting D-loop disassembly by the RecQ-type helicase Rqh1. PLoS Biol 12, e1001968.

Storici, F., and Resnick, M.A. (2006). The delitto perfetto approach to in vivo site-directed mutagenesis and chromosome rearrangements with synthetic oligonucleotides in yeast. Methods Enzymol 409, 329–345.

Sychrová, H., Chevallier, M.R., Horák, J., and Kotyk, A. (1992). Thialysine-resistant mutants and uptake of lysine in Schizosaccharomyces pombe. Curr Genet 21, 351–355.

Weisman, R., Roitburg, I., Schonbrun, M., Harari, R., and Kupiec, M. (2007). Opposite effects of tor1 and tor2 on nitrogen starvation responses in fission yeast. Genetics 175, 1153–1162.

Wood, V., Gwilliam, R., Rajandream, M.A., Lyne, M., Lyne, R., Stewart, A., Sgouros, J., Peat, N., Hayles, J., Baker, S. et al. (2002). The genome sequence of Schizosaccharomyces pombe. Nature 415, 871–880.

Yang, Y., Sterling, J., Storici, F., Resnick, M.A., and Gordenin, D.A. (2008). Hypermutability of damaged single-strand DNA formed at double-strand breaks and uncapped telomeres in yeast Saccharomyces cerevisiae. PLoS Genet 4, e1000264.

Zhang, P., Du, G., Zou, H., Chen, J., Xie, G., Shi, Z., and Zhou, J. (2016). Effects of three permeases on arginine utilization in Saccharomyces cerevisiae. Sci Rep 6, 20910.

Zhang, Y., Saini, N., Sheng, Z., and Lobachev, K.S. (2013). Genome-wide screen reveals replication pathway for quasi-palindrome fragility dependent on homologous recombination. PLoS Genet 9, e1003979.

